# TRIM24 controls induction of latent HIV-1 by stimulating transcriptional elongation

**DOI:** 10.1101/2021.11.08.467700

**Authors:** Riley M. Horvath, Matthew Dahabieh, Tom Malcolm, Ivan Sadowski

## Abstract

The conserved HIV-1 LTR *cis* elements RBE1/3 bind the factor RBF2, consisting of USF1/2 and TFII-I, and are essential for reactivation of HIV-1 by T cell signaling. We determined that TFII-I recruits the tripartite motif protein TRIM24 to the LTR, and this interaction is required for efficient reactivation of HIV-1 expression in response to T cell signaling, similar to the effect of TFII-I depletion. Knockout of *TRIM24* did not affect recruitment of RNA Pol II to the LTR promoter, but inhibited transcriptional elongation, an effect that was associated with decreased RNA Pol II CTD S2 phosphorylation and impaired recruitment of CDK9 to the LTR. These results demonstrate that TFII-I promotes transcriptional elongation in response to T cell activation through recruitment of the co-factor TRIM24, which is necessary for efficient recruitment of the elongation factor P-TEFb.

## Introduction

Despite over three decades of intensive research, human immunodeficiency virus type one (HIV-1) and acquired immunodeficiency syndrome (AIDS) remains a significant burden to human health globally, as nearly 40 million people are currently infected with this virus. Advances in development of antiretroviral therapy (ART) has eliminated HIV/AIDS as a leading cause of death globally. In most instances ART reduces plasma viremia levels to undetectable levels, but interruption of treatment allows immediate viral rebound (Laskey & Siliciano, 2014) as a consequence of the extremely long-lived viral reservoir of latently infected CD4^+^ T cells that is established early upon infection, is unaffected by ART, and escapes immune system clearance (Chun e*t al*, 1997) (Finzi *et al*, 1997) (Wong *et al*, 1997) (Joos *et al*, 2008). Various therapeutic strategies to eliminate latently infected CD4^+^ T cells, including “shock and kill” and “block and lock” are a current significant focus, which are intended to therapeutically modulate expression of the latent proviral reservoir by employing latency reversing (LRAs) or latency promoting agents (LPAs) (Sadowski & Hashemi, 2019).

HIV-1 transcription is controlled by the 5’ long terminal repeat (LTR) which serves as the viral promoter and enhancer, and contains numerous *cis* elements for host cell transcription factors (Pereira, 2000) (Sadowski *et al*, 2008). HIV-1 transcription is tightly linked to T cell activation, and consequently the LTR enhancer region contains binding sites for transcription factors that are regulated downstream of T cell activation pathways, which links viral expression to CD4^+^ T cell receptor engagement (Brooks *et al*, 2003). The Ras-response factor binding elements (RBE3 and RBE1) were initially identified as required for response of the HIV-1 LTR to activated Ras and MAPK signaling (Bell & Sadowski, 1996), and also were found to be amongst the most highly conserved LTR *cis* elements in virus from patients that develop AIDS (Estable *et al*, 1996). RBE3 (ACTGCTGA) and RBE1 (CAGCTG) are positioned at -129 and - 21 respectively, flanking the nucleosome depleted region formed by the phased nucleosomes Nuc-0 and Nuc-1 (Verdin *et al*, 1993) (Estable *et al*, 1996). These conserved elements bind a factor designated Ras-response element binding factor 2 (RBF-2), which is minimally comprised of the basic helix-loop-helix leucine zipper (b-HLH-LZ) transcription factors Upstream Stimulatory Factor 1 and 2 (USF1/2), and TFII-I (GTF2I) (Estable *et al*, 1999) (Chen *et al*, 2005). Mutation of the RBE1 and 3 elements prevent binding of RBF-2 to the LTR *in vivo* and prevents induction of HIV-1 provirus in response to T cell activation (Chen *et al*, 2005) (Malcolm *et al*, 2007) (Malcolm *et al*, 2008), although the mechanism(s) for these effects have not been elucidated. RBF-2 is also associated with YY1 at the upstream RBE3 element, and these factors may mutually regulate DNA binding to this region of the LTR (Malcolm e*t al*, 2007). However, unlike TFII-I, USF1, and USF2 which are constitutively bound to the LTR *in vivo*, YY1 is dissociated in response to T cell activation and is associated with production of immediate latent provirus which is produced within 24 hours post infection (Bernhard *et al*, 2013).

Tripartite-Motif containing protein 24 (TRIM24) possesses amino-terminal RBCC (RING, BBox, coiled-coil) domains, consistent with additional members of the TRIM protein family, in addition to a Transcription Intermediary Family (TIF1)-defining carboxy-terminal tandem plant homeodomain - bromodomain (PHD -BRD) motif (Appikonda *et al*, 2016) (Hatakeyama, 2017). *TRIM24* has been associated with both oncogenic and tumor suppressive effects, dependent on context; deletion of *TRIM24* promotes hepatocellular carcinoma (HCC) in mice, while *TRIM24* expression negatively correlates with cancer progression and patient survival rates for multiple cancers including breast and prostate (Tsai *et al*, 2010) (Herquel *et al*, 2011) (Groner *et al*, 2016). Although the mechanisms for development and progression of these cancers are ill-defined, it is thought that TRIM24 dependent transcription is involved. Specifically, TRIM24 was characterized as a cofactor for various nuclear receptors including estrogen (ER) and androgen (AR), and it was proposed that recruitment of TRIM24 to genes regulated by these factors promotes cellular proliferation and tumor growth (Tsai *et al*, 2010) (Tisserand *et al*, 2011) (Pathiraja *et al*, 2015) (Groner *et al*, 2016). TRIM24 is a co-activator of DNA sequence specific transcription factors, and this factor interacts with H3K4me0/K23ac modified histones through its tandem PHD-BRD domains (Tsai *et al*, 2010) (Groner *et al*, 2016) (Lv *et al*, 2017). Interaction of TRIM24 with chromatin promotes SUMOylation, which causes global alterations in transcription (Appikonda *et al*, 2018), but overall the molecular mechanism(s) by which this protein functions as a transcriptional co-activator has not been established. TRIM24 also has E3 ubiquitin ligase activity, and was shown to cause proteasomal mediated degradation of p53 (Allton *et al*, 2009) (Jain e*t al*, 2014). Consequently, TRIM24 may regulate cancer progression through effects on genome stability in addition to alterations in global transcriptional regulation (McAvera & Crawford, 2020).

USF1, USF2, and TFII-I are ubiquitously expressed factors that exert negative or positive effects on transcription, dependent upon promoter context and cell differentiation state (Corre & Galibert, 2005) (Roy, 2012). TFII-I in particular plays a vital role for RBF-2 formation and function as USF1 and 2 have low affinity for RBE3 and RBE1 in its absence *in vitro* (Chen *et al*, 2005) (Dahabieh *et al*, 2011). TFII-I was initially identified bound to the adenovirus major late (AdML) core promoter and transcriptional initiator (Inr) elements (Roy *et al*, 1991), but was subsequently observed associated with various sequence-specific transcription factors, including SRF, STAT1, STAT3, ATF6 and USF1/2 (Roy, 2012). TFII-I protein has six directly reiterated putative (helix-loop-helix) HLH motifs, termed I-repeats (Doi-Katayama *et al*. 2007) that may facilitate protein - protein interactions or binding to specific DNA *cis*-elements in cooperation with additional factors (Vullhorst & Buonanno, 2005). Many of these interactions promote activation of transcription, but TFII-I was also shown to cause transcriptional repression through recruitment of HDAC3 and the PRC2 component SUZ12 (Wen *et al*, 2003). Accordingly, mutation of the RBE1 and 3 elements inhibit reactivation of latent HIV-1 in response to T cell activation, but also cause elevated basal expression of LTR reporter genes (Chen *et al*, 2005) (Malcolm *et al*, 2008). Furthermore, a dominant negative TFII-I mutant also impaired reactivation of HIV-1 LTR expression in response to PMA or T-cell receptor cross linking (Chen *et al*, 2005). These observations indicate a requirement of TFII-I for regulation of HIV-1 transcription, but the precise mechanistic role for its effect on viral transcription has not been determined.

Here we provide further evidence that TFII-I is essential for induction of latent HIV-1 in response to T cell activation, and demonstrate a novel interaction between TFII-I and TRIM24. We show that TFII-I recruits TRIM24 to the HIV-1 LTR, and that loss of TRIM24 expression produces similar effects as TFII-I depletion. Furthermore, TRIM24 was found to promote efficient transcriptional elongation from the viral promoter in response to T cell signaling. These observations provide novel mechanistic insight into the function of TFII-I and regulation of HIV-1 transcription for control of reactivation from provirus latency and reveals a novel function of TRIM24 as a transcriptional coactivator.

## Results

### TFII-I is required for induction of HIV-1 expression

We have previously shown that mutation of the RBE1 and 3 binding sites for RBF-2/ TFII-I on the LTR prevents induction of HIV-1 provirus expression in response to T cell signaling. However, these elements are also necessary for binding of USF1, USF2, and YY1 (Bernhard *et al*, 2013). To examine the effect of TFII-I in isolation on LTR-directed transcription we knocked down its expression using shRNA (Figure 1B) in a Jurkat cell line bearing an HIV-1 mini-virus where luciferase is expressed from the 5’ LTR (Figure 1A). We found that knockdown of TFII-I expression significantly impaired response of LTR-directed luciferase expression in cells treated with PMA (Figure 1C). The effect of TFII-I knockdown was observed as early as 2 hours post stimulation with a combination of PMA and ionomycin, an effect that became more pronounced after 6 hours treatment (Figure 1D). These results are consistent with previous observations that mutation of the RBE *cis*-elements, which bind TFII-I, impairs response of the 5’ LTR to T cell activation signals.

**Figure 1.**
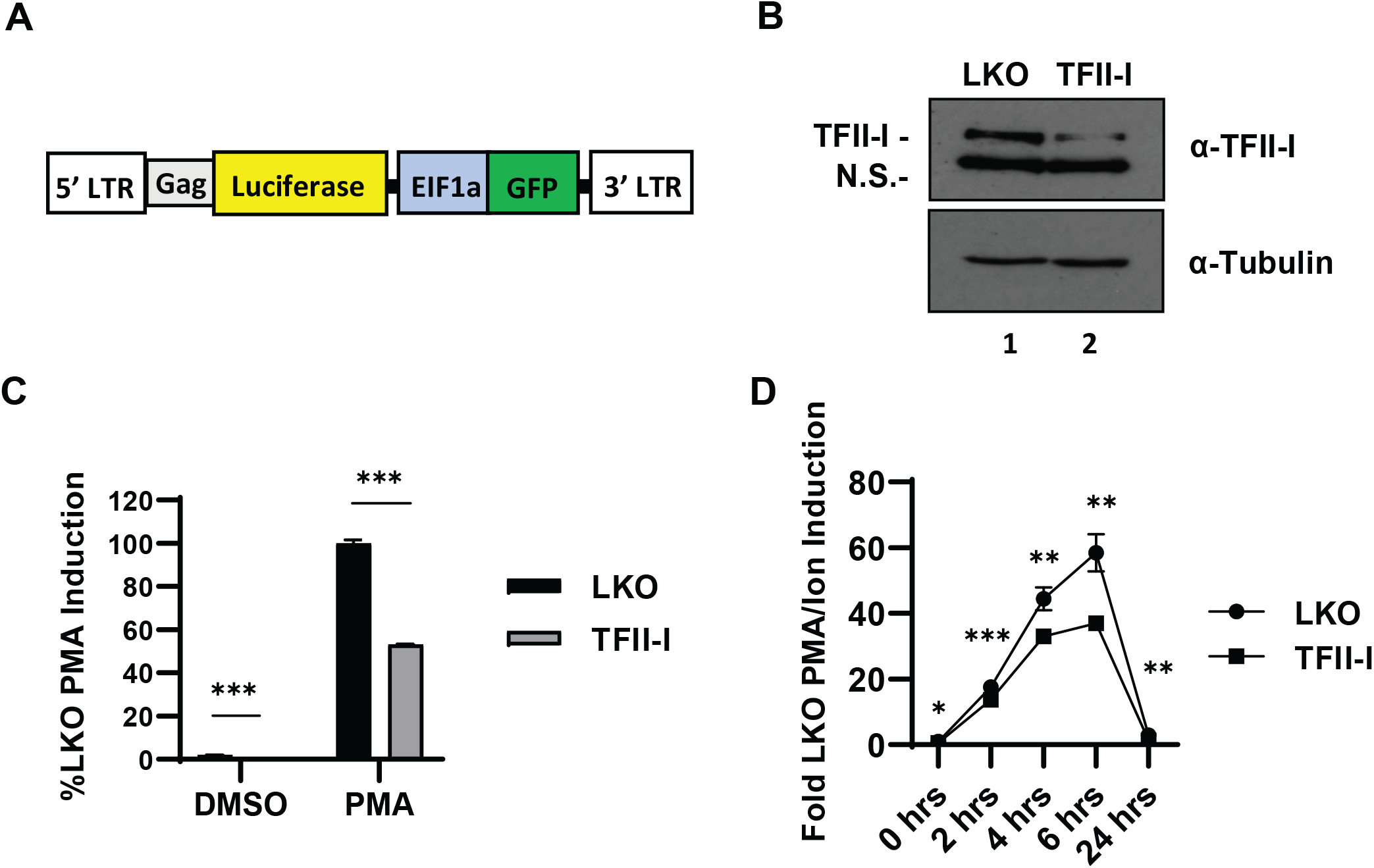
TFII-I promotes HIV-1 expression. **Panel A:** Schematic representation of HIV-1 mini-virus in the KB001 Jurkat Tat cell line, where Luciferase is expressed from the 5’ LTR as a fusion with Gag. **Panel B:** KB001 cells were transduced with pLKO lentivirus vector control (LKO, lane 1) or expressing TFII-I shRNA (lane 2). Immunoblots were performed on whole cell lysates prepared 8 days post selection in puromycin, with antibodies against TFII-I (top) or tubulin (bottom). **Panel C:** KB001 cells transduced with LKO vector control or expressing TFII-I shRNA were left untreated (DMSO) or stimulated with 20 nM PMA for 4 hours prior to measuring luciferase activity, assays were performed in triplicate and error bars represent standard deviations. **Panel D:** Transduced KB001 were treated with 20 nM PMA/ 1 µM ionomycin for the indicated time prior to measuring luciferase activity. Results are an average of three determinations, and error bars represent standard deviations.

### TFII-I interacts with TRIM24 in vivo

To identify mechanisms for regulation of HIV-1 transcription by TFII-I, we looked for novel interacting proteins using a 2-hybrid screen. Because TFII-I causes activation of transcription in yeast when fused to a DNA-binding domain, we used a modified 2-hybrid strategy designed for transcriptional activators where interaction between bait and prey fusions causes repression of transcription by recruitment of the repressor protein TUP1 (Hirst *et al*, 2001). Using a GAL4-TFII-I bait protein we isolated multiple clones encoding TUP1-TRIM24 fusions from a human T cell cDNA library. To determine whether this interaction identified in yeast can be detected in human cells, we co-transfected HEK293T cells with plasmids expressing epitope tagged TFII-I and TRIM24 where we observe TRIM24-myc associated with TFII-I-Flag immunoprecipitated from cell lysates (Figure 2A, lane 6). We performed a similar experiment in the Jurkat T cell line where we expressed TRIM24-Flag and observed interaction with endogenous TFII-I by co-immunoprecipitation (Figure 2B, lane 4).

**Figure 2.**
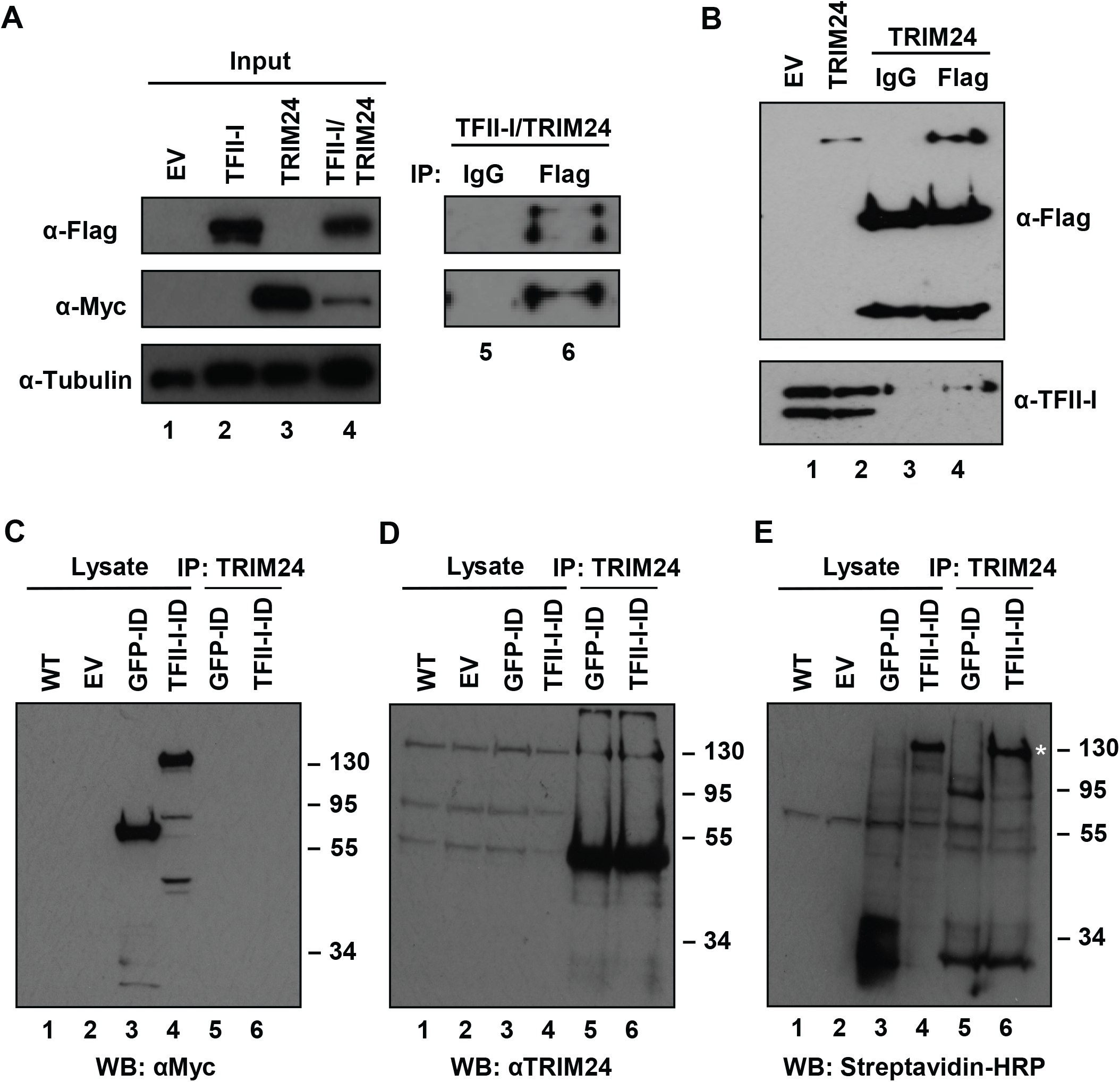
TRIM24 interacts with TFII-I *in vivo*. **Panel A:** HEK293T cells were transfected with plasmids expressing TFII-I-Flag and/ or Trim24-myc, or an empty vector. (Lanes 1 to 4) Lysates were analyzed by immunoblotting with antibodies against the Flag and myc epitopes, or tubulin. Cells co-expressing TFII-I-flag and Trim24-myc were immunoprecipitated with control (IgG) (lane 5) or Flag (lane 6) antibodies and complexes analyzed by immunoblotting with Flag and myc antibodies as indicated. **Panel B:** Lysates from Jurkat cells bearing a vector control (EV, lane 1) or expressing TRIM24-Flag (lane 2) were analyzed by immunoblotting with antibodies against Flag peptide (top) or TFII-I (bottom). Jurkat cells expressing TRIM24-Flag were immunopreciptated with control (IgG, lane 3), or anti-Flag (lane 4) antibodies, and complexes analyzed by immunoblotting with anti-Flag or anti-TFII-I antibodies as indicated. **Panels C, D and E:** HEK293T (lane 1) or cells expressing GFP-TurboID-myc (lane 3 and 5), TFII-I-TurboID-myc (lane 4 and 6), or an empty vector (lane 2) were incubated with 500 µM biotin for 1 hr. Cell lysates (lanes 1-4) or TRIM24 immunoprecipitates (lanes 5-6) were analyzed by immunoblotting with antibodies against myc (**Panel C**), TRIM24 (**Panel D**), or with Streptavidin-HRP (**Panel E**). Biotinylated TRIM24 is indicated with a white asterisk (Panel E, lane 6).

We also examined interaction of TFII-I and TRIM24 in HEK293T cells using proximity labeling with Bio-ID technology. For this analysis, we transfected HEK293T cells with plasmids expressing TFII-I or GFP as fusions with BirA^*^ (TurboID) and myc epitope tags (Figure EV1A). Upon treatment with biotin, we observed that the TFII-I and GFP TurboID fusions cause biotinylation of cellular proteins as early as 1 hour post treatment, and this effect persisted up to 24 hours (Figure EV1B). Importantly, we found that TRIM24 was biotinylated in cells expressing the TFII-I-BirA^*^ fusion (Figure 2E, lane 6), but not in cells expressing GFP-BirA^*^ (lane 5). Proximity labeling of TRIM24 by TFII-I-BirA^*^ indicates that these factors associate in the context of living cells and supports the contention that TFII-I and TRIM24 interact *in vivo*.

### TRIM24 activates expression from the HIV-1 LTR

Because TRIM24 is capable of interaction with TFII-I in both HEK293T and Jurkat cells, we next examined whether this factor affected expression from HIV-1 LTR. We co-transfected HEK293T cells with an LTR-luciferase reporter and varying amounts of plasmid expressing TRIM24-myc or TFII-I-Flag (Figure 3A). We observed a dose dependent, three-fold increase in transcriptional activity of LTR-Luc relative to endogenous levels of TRIM24 (Figure 3B). In contrast, we observed only a modest 1.3 fold increase in transcription when TFII-I-Flag was overexpressed. These results are consistent with the previously described role of TFII-I at the HIV-1 LTR (Chen *et al*, 2005) (Dahabieh *et al*, 2011), and a model of TFII-I mediated TRIM24 recruitment, where TRIM24 may represent a limiting factor for activation of HIV-1 transcription.

**Figure 3.**
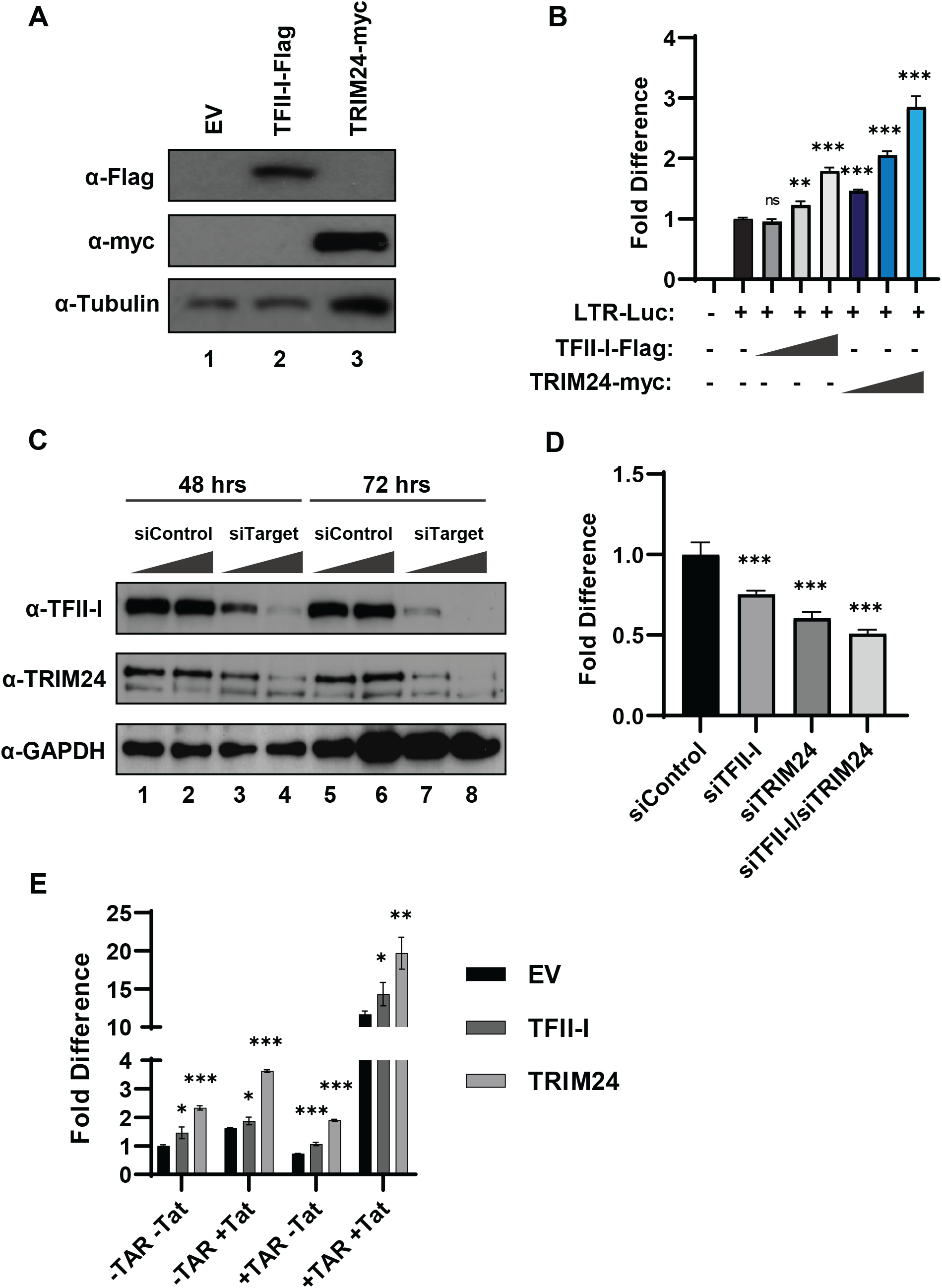
HIV-1 LTR transcription is activated by TRIM24. **Panel A:** HEK293T cells were transfected with plasmids expressing TFII-I-Flag (lane 2), TRIM24-myc (lane 3), or an empty vector (EV, lane 1). Lysates were analyzed by immunoblotting against myc, Flag, or tubulin as indicated. **Panel B:** HEK293T cells co-transfected with an LTR-Luciferase reporter plasmid and TFII-I-Flag or TRIM24-myc expression vectors were analyzed for Luciferase activity. Results are an average of three independent transfections, and error bars represent standard deviation. **Panel C:** HEK293T cells were transfected with siRNA against TFII-I or TRIM24 (lanes 3-4 and lanes 7-8). Lysates were collected 2-(lanes 1-4) and 3-days (lanes 5-8) post transfection and analyzed by immunoblotting with anti-TRIM24, anti-TFII-I, or anti-GAPDH antibodies as indicated. **Panel D:** HEK293T cells co-transfected with an LTR-Luciferase reporter plasmid and siRNA targeting TFII-I and/or TRIM24 were analyzed for Luciferase expression. Assays were performed in triplicate and error bars represent standard deviations. **Panel E:** HEK293T cells were co-transfected with an LTR-luciferase reporter plasmid and TRIM24-myc and/or TFII-I-Flag expression vector, in the presence (+ Tat) or absence (-Tat) of Tat expression vector. The LTR-luciferase reporter contained (+TAR) or lacked the TAR (-TAR) stem loop region. Results are the average of three measurements and error bars represent standard deviations.

To further examine the role of TRIM24 for transcriptional activation of HIV-1 expression, we determined the effects of TRIM24 and TFII-I knockdown. We first validated siRNA SmartPools (Dharmacon) against TRIM24 and TFII-I in transiently transfected HEK293T cells. Both siRNA pools were efficient at knocking down their target factors as determined by immunoblotting, as early as 48 hours post transfection; however, knockdown was more substantial at 72 hours post transfection (Figure 3C). We co-transfected the siRNA pools with an LTR-luciferase reporter construct into HEK293T cells, and measured luciferase activity 72 hours post transfection. Here we observed a 40% decrease in luciferase activity in TRIM24 knockdown cells, relative to control cells (Figure 3D). Consistent with results from overexpression, knockdown of TFII-I resulted in a more modest 25% decrease in luciferase activity, and knockdown of both TFII-I and TRIM24 produced an effect similar to knockdown of TRIM24 alone (Figure 3D).

We also examined the effect of TRIM24 for Tat-dependent activation of the HIV-1 LTR, where we transiently transfected HEK293T cells with LTR-luciferase reporter constructs, either bearing or lacking the Tat-responsive element TAR, in the presence or absence of a plasmid expressing the viral transactivator Tat as previously described (Dahabieh *et al*, 2011). Regardless of the presence of TAR and Tat, overexpression of TRIM24 and TFII-I resulted in elevated expression of luciferase expression (Figure 3E), indicating that the effect of TRIM24 on LTR-directed expression is largely independent of the function of HIV-1 Tat.

### TRIM24 is required for induction of HIV-1 provirus expression

The results shown above indicate that TRIM24 interacts with TFII-I *in vivo*, and its presence correlates with activation of HIV-1 LTR-directed transcription in HEK293T cells. We next examined the requirement of this factor for HIV-1 provirus expression. For this, we produced CRISPR-Cas9 mediated gene disruption in a Jurkat T cell line (Figure 4A) bearing a chromosomally integrated HIV-1 mini-virus in which luciferase expression is driven by the 5’ LTR (Figure 1A). Interestingly, for each of these *TRIM24* knockout (KO, *trim24* KO) cell lines, we observed a significant reduction in HIV-1 expression in response to treatment with PMA, as measured by luciferase activity (Figure 4B). We observed a similar effect using shRNA mediated *TRIM24* knockdown (Figure EV2).

**Figure 4.**
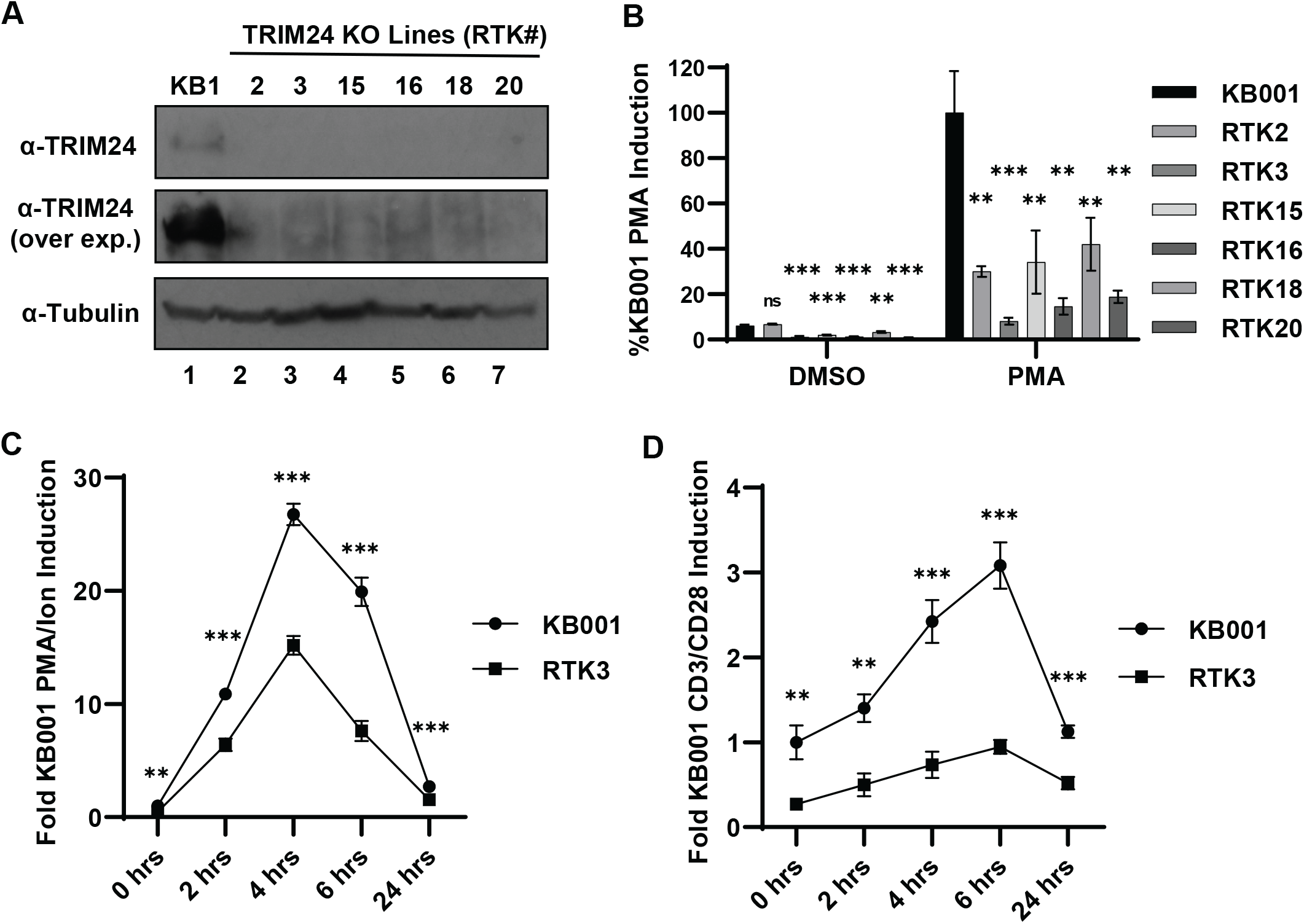
TRIM24 is required activation of HIV-1 in T cells. **Panel A:** Knockout of *trim24* (lanes 2-7) in Jurkat cell line KB001 (lane 1) were produced using CRISPR/ Cas9. Following expansion of clones, lysates were analyzed by immunoblotting using antibodies against TRIM24 (top and center, over-exposed) or tubulin (bottom). **Panel B:** KB001 parent and RTK *trim24* KO clonal cell lines were left untreated (DMSO) or treated with 20 nM PMA for 4 hrs at which point Luciferase activity was measured. Assays were performed in triplicate and error bars represent standard deviations. **Panel C:** KB001 and RTK3 *trim24* KO cell lines were incubated with 20 nM PMA/ 1 μ M ionomycin. Luciferase expression was analyzed at the indicated time points. Results are an average of three measurements, and error bars represent standard deviations. **Panel D:** As in Panel C but cells were cultured in the presence of CD3/CD28 coated beads.

Induction of HIV-1 expression was significantly decreased in *trim24* knockout cells up to at least 6 hours post stimulation with PMA/ ionomycin (Figure 4C) or by T cell receptor cross linking (Figure 4D). These results indicate that TRIM24 is essential for rapid and full induction of HIV-1 expression in response to T cell activation signals.

### TRIM24 localizes to the HIV-1 LTR and is TFII-I dependent

Our results indicate that TFII-I and TRIM24 positively regulate HIV-1 expression and that TRIM24 may be a limiting factor for transcriptional reactivation by T cell signaling. To further assess this possibility, we examined TFII-I and TRIM24 occupancy on the HIV-1 LTR using ChIP and analysis by Q-PCR. Consistent with previous observations (Bernhard *et al*, 2013), we observe interaction of TFII-I at the LTR regions representing both RBE3 and RBE1, which is not significantly altered by treatment with PMA (Figure 5A), and a similar result was observed for TRIM24 (Figure 5B); however we do observe recruitment of NFκB p65 to the enhancer region in response to PMA treatment of these same cells (Figure EV3). These observations indicate that, consistent with the interaction as measured by co-IP and Bio-ID, TRIM24 co-localizes with TFII-I on the HIV-1 LTR *in vivo*. We also observed association of TRIM24 with the HIV-1 LTR in the HeLa-derived TZM-bl cell line, indicating that this observation is preserved among cell tissue types (Figure EV4).

**Figure 5.**
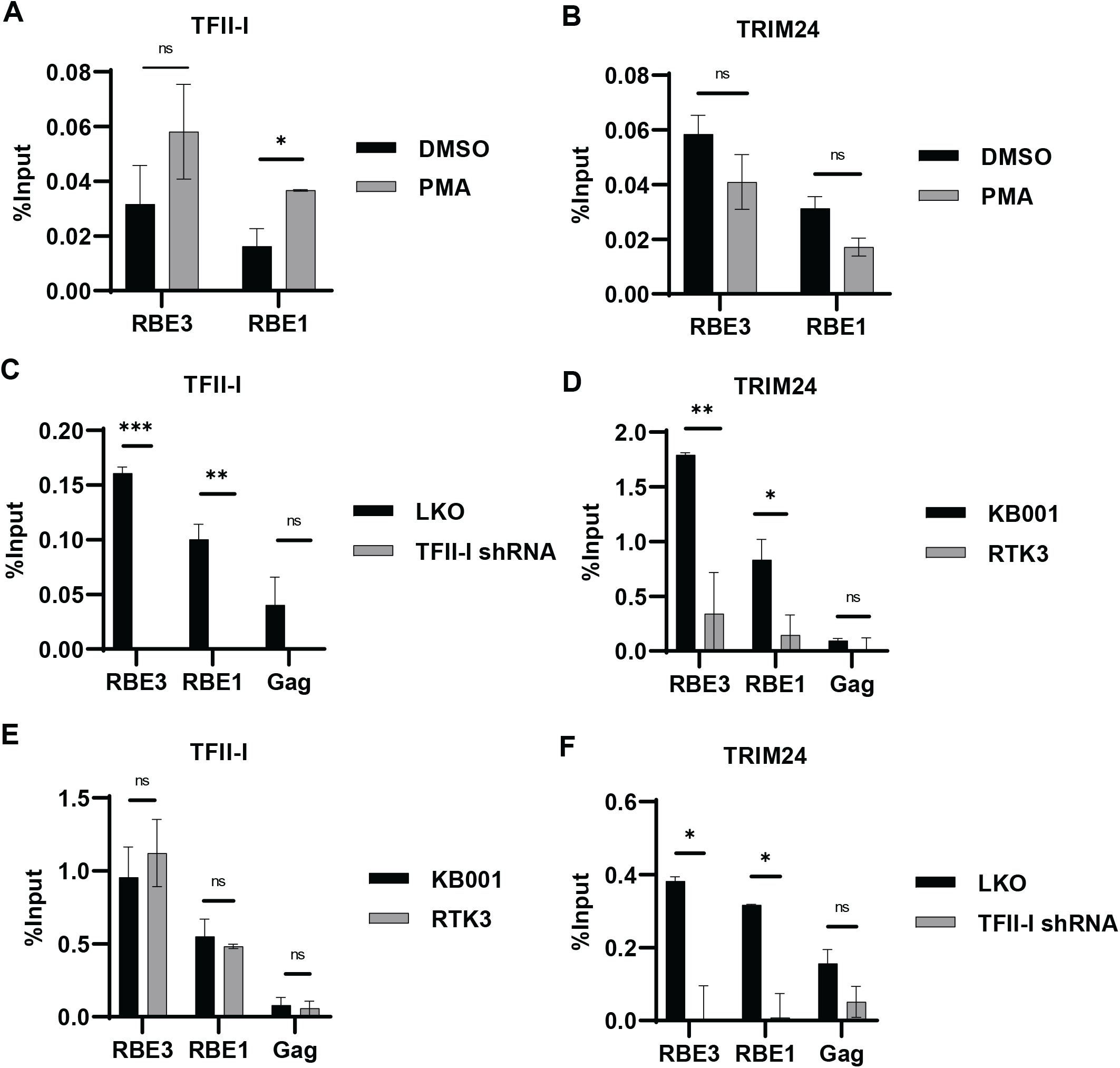
TFII-I/ TRIM24 co-localize to the HIV-1 LTR. **Panels A and B:** Jurkat Tat mdHIV clone 11 cells were left untreated (DMSO) or incubated with 50 nM PMA for 24 hrs. ChIP-qPCR was performed using anti-TFII-I (Panel A) or anti-TRIM24 (Panel B) antibodies. **Panel C:** ChIP-qPCR analysis was performed using TFII-I antibodies with KB001 cells transduced with pLKO lentiviral control vector or expressing TFII-I shRNA. **Panel D:** KB001 or RTK3 *trim24* KO cell lines were subject to ChIP-qPCR analysis using anti-TRIM24 antibodies. **Panel E:** As in Panel D but anti-TFII-I antibodies used for ChIP analysis. **Panel F:** As in Panel C but anti-TRIM24 antibodies used for ChIP analysis. All results are normalized by subtraction of values produced with non-specific IgG and represent average of ≥2 independent experiments with error bars representing standard deviations.

To examine the requirement of TFII-I for recruitment of TRIM24 to the LTR and *vice versa*, we performed ChIP-qPCR in cells where these factors were depleted. We found that interaction of TFII-I with the LTR was unaffected by the *trim24* knockout, relative to wild type cells (Figure 5E). In contrast, interaction of TRIM24 with both the RBE3 and RBE1 regions of the LTR was significantly reduced in cells where TFII-I expression was knocked down with shRNA (Figure 5F). Thus, loss of TRIM24 has no effect on binding of TFII-I to its sites on the LTR but interaction of TRIM24 with the LTR is completely dependent upon TFII-I. These observations support the contention that TFII-I directly recruits TRIM24 to the HIV-1 LTR.

### TRIM24 stimulates HIV-1 transcriptional elongation

Having observed an essential role of TFII-I and TRIM24 for induction of HIV-1 expression, we sought to identify the specific defect in HIV-1 transcriptional response caused by loss of TRIM24. To assess the impact of TRIM24 on initiation and elongation of LTR bound RNAP, we employed a previously characterized RT-PCR assay (Zhu et al., 2012) that takes advantage of the distinct mRNA sequences present in initiating versus elongating (proximal, Gag) transcripts (Figure 6A). In these experiments, we did not observe a difference in abundance of HIV-1 RNA associated with paused RNA Polymerase II between untreated WT and *TRIM24* KO cells (Figure 6B, Initiation). However, we did observe a marked decrease in proximal and Gag mRNA transcripts in *TRIM24* KO cells relative to WT untreated cells (Figure 6B), and this difference was significantly exaggerated in cells stimulated with PMA (Figure 6C), indicative of diminished RNA polymerase processivity. This result indicates that TRIM24 might facilitate elongation of RNA polymerase from the LTR core promoter.

**Figure 6.**
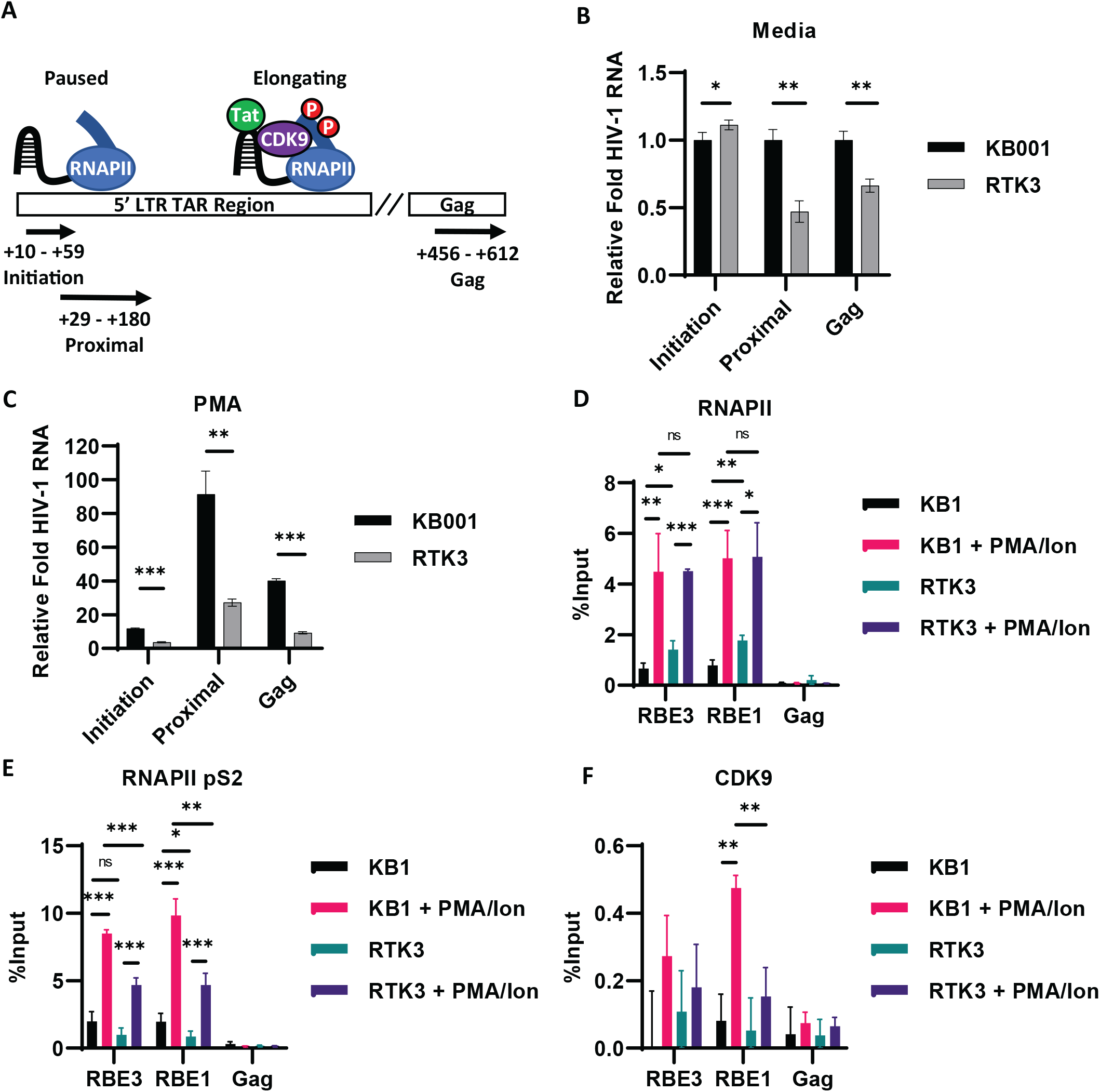
TRIM24 stimulates elongation of HIV-1 transcription. **Panel A:** Schematic representation of the 5’ LTR TAR and Gag-encoding regions, indicating primer pairs used to measure Initiating (+10-59 bp), Proximal (+29-180 bp), or Gag (+456-612 bp) HIV-1 transcript abundance. **Panels B and C:** KB001 and RTK3 *trim24* KO cells were cultured under normal conditions (Panel B) or with 20 nM PMA for 4 hrs (Panel C). RNA was extracted, cDNA synthesized, and RT-PCR was performed using primers to detect Initiating, Proximal and Gag RNA sequences. Values are an average of three replicates, and error bars represent standard deviation. Data was normalized to the untreated KB001 control. **Panels D, E, and F:** KB001 or RTK3 *trim24* KO cells were left untreated or stimulated with 20 nM PMA/ 1 μ M ionomycin for 4 hrs. ChIP-qPCR analysis was performed using anti-RNAPII (Panel D), anti-RNAPII pS2 (Panel E), or anti-CDK9 (Panel F) antibodies. Results were normalized by subtraction of values produced with non-specific IgG, and are an average of ≥3 independent ChIP experiments. Error bars represent standard deviations.

We also examined the effect of TRIM24 on recruitment of RNA Polymerase to the LTR. We observe enhanced interaction of RNA Polymerase with the LTR in cells treated with PMA and ionomcyin, relative to untreated cells, as measured by ChIP Q-PCR. However, although the *TRIM24* knockout caused a slight increase in RNA Pol II occupancy at the LTR promoter in untreated cells, relative to wild type, we did not observe a difference in cells treated with PMA and ionomycin (Figure 6D). This result indicates that TRIM24 is dispensable for initial recruitment of RNAPII to the LTR.

RNA Polymerase II becomes phosphorylated at S2 on the CTD during initiation of transcription, an effect that promotes transcriptional elongation (Buratowski, 2009). As expected, ChIP-qPCR analysis of phosphorylated Ser2 (pS2) CTD showed enrichment upon stimulation of T cell signaling in wild type cells (Figure 6E, KB1). However, accumulation of pS2 modified RNAPII was significantly reduced in the *trim24* knockout cell line (Figure 6E, RTK3). Phosphorylation of CTD S2 is produced by CDK9 of P-TEFb (Bieniasz *et al*, 1999), and consequently we also examined whether the *TRIM24* knockout affected recruitment of CDK9 to the LTR. In agreement with observations that P-TEFb is sequestered by the 7SK snRNP complex in unstimulated T cells (Mbonye *et al*, 2013), ChIP-qPCR using antibodies against CDK9 revealed low levels of association at the LTR in untreated cells, but significant enhancement in WT cells treated with PMA and ionomycin (Figure 6F, KB1). In contrast, we did not observe a significant increase of CDK9 occupancy in stimulated *TRIM24* depleted cells (Figure 6F, RTK3). Importantly, we do not observe an effect of the *TRIM24* knockout on expression of CDK9 protein, or cyclin T1 (Figure EV5). Consistent with previous observations (Mbonye *et al*, 2018), we find that treatment of cells with a CDK9 inhibitor prevented reactivation of latent HIV-1 in response to treatment with PMA (Figure EV6). Collectively, these results indicate that TRIM24 mediates activation of HIV-1 expression by promoting recruitment of CDK9 to the viral promoter and facilitating the transition of RNAPII into an actively elongating complex.

## Discussion

In this study we identified a novel interaction between TFII-I and TRIM24, which we found to be required for efficient transcriptional elongation from the HIV-1 LTR promoter. The important role of transcriptional elongation for regulation of HIV-1 expression was recognized since discovery that the viral transactivator Tat recruits P-TEFb to paused RNA Polymerase II complexes through interaction with nascent TAR RNA (Bieniasz *et al*, 1999). Accordingly, loss of Tat in cells bearing transcriptionally silenced provirus is thought to represent a significant barrier for reactivation of virus expression (Karn, 2011). Interestingly however, one report has indicated that several commonly studied latency reversing agents (LRA), including PEP005 and Ingenol cause reactivation of viral expression by promoting transcriptional elongation and splicing of sub-genomic transcripts, rather than initiation of transcription (Yukl *et al*, 2018), although mechanism(s) for this effect were not determined. Our results indicate that Tat or TAR are not required for stimulation of HIV-1 transcription by TRIM24. Consequently, it is possible that the effect of TFII-I-TRIM24 for recruitment of CDK9, and phosphorylation of CTD S2, might represent a priming mechanism to kick start production of elongated and spliced transcripts for production of Tat protein (Mbonye & Karn, 2014).

The function of TFII-I for regulation of HIV-1 transcription has been enigmatic. Binding sites for TFII-I, in conjunction with USF1 and 2 are highly conserved on LTRs from patients, and these elements are necessary for reactivation of HIV-1 transcription in response to T cell signaling (Chen *et al*, 2005). This observation is consistent with results indicating that TFII-I is involved in activation of *c-Fos* expression in response to MAPK signaling (Kim & Cochran, 2000), but a specific mechanistic role of TFII-I for stimulation of transcription from the HIV-1 LTR has not previously been identified. TFII-I interacts with multiple additional DNA binding factors, including STAT1, STAT3, SRF, ERSF, and on the HIV-1 LTR USF1 and USF2 (Roy, 2012) (Chen *et al*, 2005). In the latter case, interaction of USF1/2 with the upstream RBE3 element on the HIV-1 LTR requires interaction with TFII-1 to produce a specificity unique to this complex (Malcolm *et al*, 2008). These observations suggest that TFII-I plays a significant role for directing cooperative interaction of various sequence specific factors with regulatory *cis*-elements. However, our finding that TFII-I recruits the co-activator TRIM24 suggests this factor plays a more elaborate role for gene regulation than acting as a chaperone for DNA binding partners. It will be interesting to determine how additional functions attributed to TRIM24 are separated from those involved in interaction with TFII-I and recruitment to the HIV-1 LTR.

TRIM24 was previously identified as a co-activator for various nuclear hormone receptors, including for estrogen (Thénot *et al*, 1997), androgen (Kikuchi *et al*, 2009), and androstane (Kanno *et al*, 2018). However, the mechanism(s) by which TRIM24 causes transcriptional activation by nuclear receptors has not been identified. Considering our results, we propose that recruitment of TRIM24 to target genes of these receptors may also cause activation of transcriptional elongation. Results shown here indicate that TRIM24 is required for efficient recruitment of CDK9 to the HIV-1 LTR. We imagine various functions of TRIM24 that might contribute to this effect, the simplest hypothesis being that TRIM24 may directly interact with CDK9 or cyclin T to promote recruitment of P-TEFb (Figure 7). Alternatively, TRIM24 may modify additional factors at the HIV-1 LTR core promoter to enable access or recruitment of P-TEFb. TRIM24 has E3-ubiquitin ligase activity, and was shown to promote degradation of p53 through direct ubiquitylation (Allton *et al*, 2009) (Jain *et al*, 2014). TRIM24 was also found to modify the functions of CBP and TRAF3 by catalyzing K63-linked ubiquitination. TRIM24 mediated K63 ubiquitination of CBP inhibited macrophage polarization (Yu *et al*, 2019), while this modification of TRAF3 was observed as part of the anti-viral response to VSV infection (Zhu *et al*, 2020). Consequently, it is possible that the role of TRIM24 for recruitment of P-TEFb to the HIV-1 promoter may involve post-translational modification of one or more target factors that promote recruitment of P-TEFb to the LTR promoter. Elucidation of the precise role of TRIM24 as a transcriptional co-activator will require a detailed understanding of its interactions with the general transcription factor machinery. Nevertheless, our results provide a novel glimpse towards understanding the function of TRIM24 as a transcriptional co-activator.

**Figure 7.**
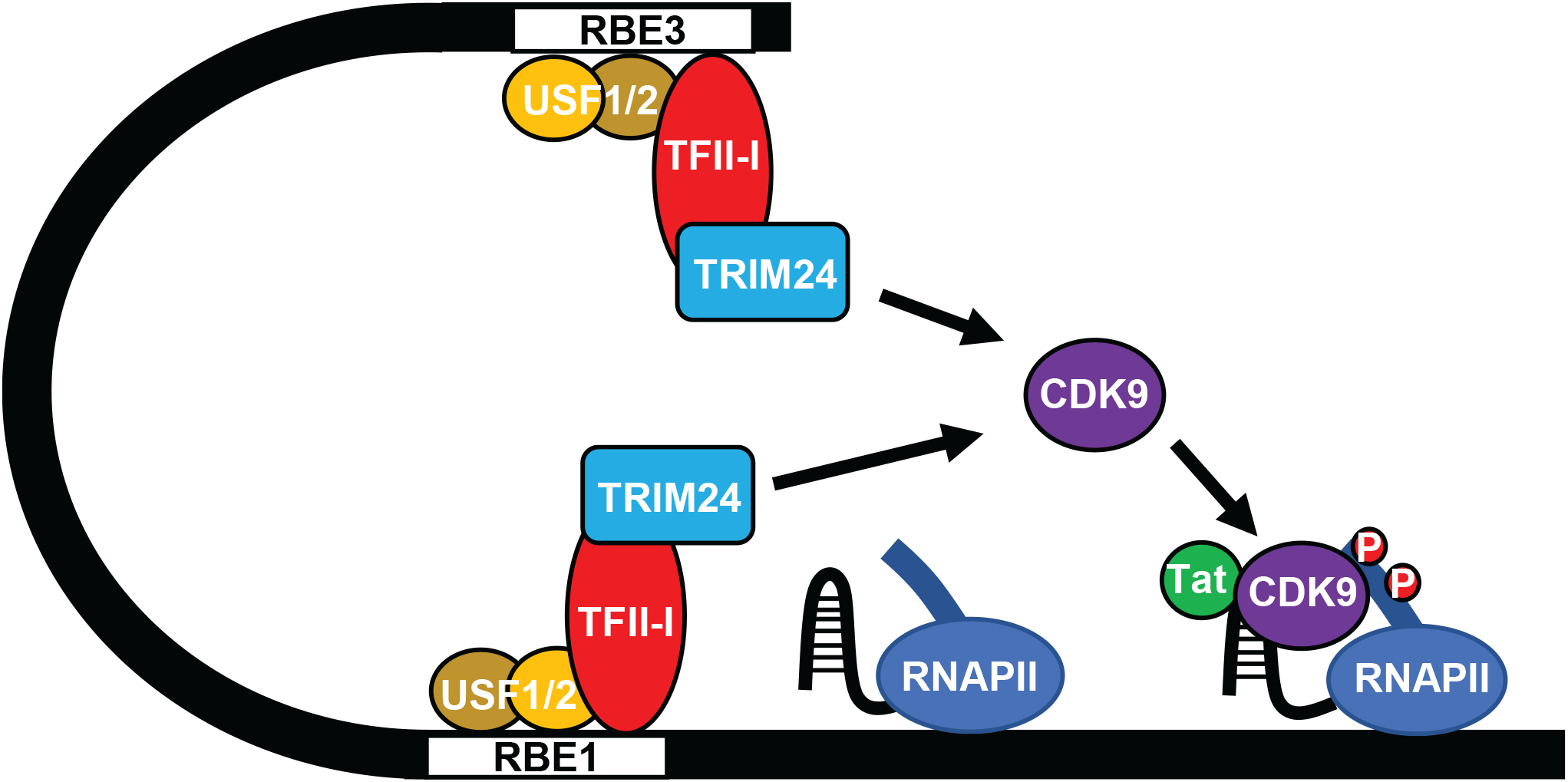
TRIM24 is an RBF-2 cofactor that stimulates transcriptional elongation of HIV-1. The TFII-I component of RBF-2 directly interacts with, and recruits TRIM24 to the HIV-1 LTR. TRIM24 is essential for T cell signal induced activation of chromosomally integrated provirus and is required for recruitment of P-TEFb/CDK9 to the LTR, and phosphorylation of serine 2 of the RNA Pol II CTD, to promote transcriptional elongation.

There is currently considerable interest in development of therapies that could be applied in addition to or instead of antiretroviral therapy to eliminate cells latently infected with HIV-1 from patients. One potential strategy, broadly termed “block and lock” is based on the rationale that preventing stochastic basal transcriptional noise of the latent provirus may discourage maintenance of the latently infected cell population (Sadowski & Hashemi, 2019). Based on results presented here, we suggest that TRIM24, and specifically the interaction between TRIM24 and TFII-I may represent an important specific target for this strategy. Consequently, a more detailed understanding of the interaction between these factors will be important for development of this possibility.

## Materials and Methods

### Cell and virus culture

Jurkat E6-1, Jurkat Tat, and TZM-bl cells were cultured under standard conditions as previously described (Dahabieh *et al*, 2011). Vesicular stomatitis virus G (VSV-G) pseudotyped viral stocks were produced by co-transfecting HEK293T cells with a combination of viral molecular clone, psPAX, and pHEF-VSVg as previously described (Dahabieh *et al*, 2011).

### shRNA knockdown

Jurkat cells were infected with pLKO empty vector (Addgene #8453) or pLKO shRNA expressing lentivirus at a M.O.I. ∼10. TFII-I shRNA infected cells were cultured for 8 days with 7.5 µg/mL puromycin while TRIM24 shRNA infected cells were cultured 3 days with 3 µg/mL puromycin. Pools of puromycin selected cells were prepared for the indicated analysis. MISSION shRNA clones (Sigma) in pLKO.1 backbone were: TFII-I, TRCN0000019315; TRIM24 A, TRCN0000021263; TRIM24 B, TRCN0000021262; TRIM24 C, TRCN0000021259.

### TRIM24 knockout

*TRIM24* knockout cell lines were generated using CRISPR-Cas9 in a Jurkat Tat cell line possessing chromosomally integrated HIV-1 mini-virus where luciferase is expressed from the 5’ LTR (Figure 1A). Briefly, 2×10^6^ cells were co-transfected with Cas9 (pU6_CBh-Cas9-T2A-BFP: Addgene #64323) and gRNA (pSPgRNA: Addgene #47108) sequences that target genomic *TRIM24*, using the Neon Transfection System (Invitrogen) as per the manufacturer’s instructions. Knockout cells were isolated by live sorting (Astrios Flow Cytometer) BFP positive cells into 96-well plates containing complete RPMI 1640. Clones were expanded, and *TRIM24* KO was validated by PCR genotyping and western blotting. *TRIM24* gRNA target sequences were CTGCATATTATTTAAGCAAC and GAACGAGGCCGAGAGTCGGC.

### Immunoblotting and Immunoprecipitation

Western blotting was performed as previously described (Hashemi *et al*, 2018). Antibodies were as follows: Tubulin - Abcam #ab7291, TFII-I - Abcam #ab134133, Flag - Sigma Aldrich #F3165, Myc - Santa Cruz #sc-40, TRIM24 - Proteintech #14208-1-AP, GAPDH - Abcam #ab9484, CDK9 - Abcam #ab239364, Cyclin T1 - Santa Cruz #sc-10750, Streptavidin-HRP - Abcam #ab7403, Goat Anti-Rabbit-HRP - Abcam #ab6721, Goat Anti-Mouse-HRP - Pierce #1858413.

Immunoprecipitations (IPs) were performed using HEK293T or Jurkat E6-1 cells. 8.33×10^5^ HEK293T cells were plated with 2 mL DMEM in 6-well plates. The following day, cells were transfected with 4 μg of the indicated construct, using 3 µg PEI per 1 µg plasmid DNA (Durocher *et al*, 2002) and harvested 24-48 hrs post-transfection. For Jurkat IPs, cells were transduced with TRIM24-Flag or empty vector lentivirus and experiments were performed following puromycin selection. Cells were collected, washed with ice cold PBS, suspended in DR Buffer A (10 mM HEPES-KOH pH = 7.9, 10 mM KCl, 1.5 mM MgCl_2_, 0.5 mM DTT, 1x PIC, 0.5 mM PMSF) and incubated at 4^0^C for 15 min. Samples were centrifuged, the cytoplasmic supernatant was discarded while the pelleted nuclei were suspended in DR Buffer C (20 mM HEPES-KOH pH = 7.9, 0.42 M NaCl, 1.5 mM MgCl_2_, 0.2 mM EDTA, 25% glycerol, 0.5 mM DTT, 1x PIC, 0.5 mM PMSF) and briefly sonicated (Covaris S220 Focused-ultrasonicator). Samples were cleared by centrifugation and protein concentrations were determined using a Bradford assay (BioRad). 250 μg of nuclear extract was diluted with Flag-IP Buffer (50 mM Tris-HCl pH = 8.0, 90 mM NaCl, 1 mM EDTA, 1% Triton X-100, 1x PIC) and antibody was added (Flag - Sigma Aldrich #F3165, mouse IgG – Santa Cruz #sc-2025); the antibody – lysate mixture was incubated at 4°C with rotation for 1 hr prior to the addition of protein A/G agarose beads (Millipore, 50 μL/IP) and continued overnight incubation. The samples were washed 3x with Flag-IP Buffer and eluted in 4x SDS Sample Buffer with boiling. Eluted materials were subject to analysis by immunoblotting.

### Chromatin Immunoprecipitation

Exponentially growing Jurkat Tat KB001 or mdHIV clone 11 cells (3×10^7^ cells/IP) were fixed with 1% formaldehyde (Sigma-Aldrich) for 10 min at room temperature. Cross-linking was quenched with 125 mM glycine for 5 min, at which point cells were collected and washed with ice cold PBS. Cells were incubated in NP-40 Lysis Buffer (0.5% NP-40, 10 mM Tris-HCl pH = 7.8, 3 mM MgCl_2_, 1x PIC, 2.5 mM PMSF) for 15 min on ice. Following sedimentation, supernatant was discarded, and the pellet was resuspended in Sonication Buffer (10 mM Tris-HCl pH = 7.8, 10 mM EDTA, 0.5% SDS, 1x PIC, 2.5 mM PMSF). Nuclei were sonicated using a Covaris S220 Focused-ultrasonicator to produce sheared DNA between 2000-200 bp. Samples were pelleted, with the soluble supernatant collected as the chromatin fraction and snap frozen in liquid nitrogen. Chromatin concentrations were normalized among samples and pre-cleared with Protein A/G agarose (Millipore, 100 μL/IP). Following dilution in IP buffer (10 mM Tris-HCl pH = 8.0, 1.0% Triton X-100, 0.1% Deoxycholate, 0.1% SDS, 90 mM NaCl, 2 mM EDTA, 1x PIC), antibodies were added (TFII-I - BD Biosciences #610842, TRIM24 - Proteintech #14208-1-AP, RNAPII - Abcam #ab26721, RNAPII pS2 - Abcam #ab238146, CDK9 - Abcam #ab239364, NFκB p65 - Thermo Fisher #51-0500) and the chromatin/ antibody mixture was incubated 1 hr at 4°C with rotation. Pre-washed Protein A/G agarose beads (40 μL/IP) were then added to the samples and incubated overnight at 4°C with rotation. Bead – antibody complexes were washed 3x in Low Salt Wash Buffer (20 mM Tris-HCl pH = 8.0, 0.1% SDS, 1.0% Triton X-100, 2 mM EDTA, 150 mM NaCl, 1x PIC) and 1x with High Salt Wash Buffer (same but with 500 mM NaCl). Elution and crosslink reversal was performed by incubating 4 hrs at 65^0^C in EB supplemented with RNase A. DNA was purified using the QIAQuick PCR purification kit (QIAGEN) and ChIP DNA was analyzed using the Quant Studio 3 Real-Time PCR system (Applied Biosystems). Oligos used for ChIP-qPCR are as follows; RBE3: Fwd 5’ AGCCGCCTAGCATTTCATC, Rev 5’ CAGCGGAAAGTCCCTTGTAG. RBE1: Fwd 5’ AGTGGCGAGCCCTCAGAT, Rev 5’ AGAGCTCCCAGGCTCAAATC. Gag: Fwd 5’ AGCAGCCATGCAAATGTTA, Rev 5’ AGAGAACCAAGGGGAAGTGA. NFκB Enhancer Region (NER): Fwd 5’ TTTCCGCTGGGGACTTTC, Rev 5’ CCAGTACAGGCAAAAAGCAG.

### BioID assays

8.33×10^6^ HEK293T cells were plated in 2 mL DMEM in 6-well plates and incubated overnight. Cells were transfected using PEI with 3 μg of the indicated TurboID construct and incubated overnight. Biotin was added directly to the media at a final concentration of 500 μM and cells were incubated for 1 hr. Biotinylation was stopped by washing cells 2x in ice cold PBS. Endogenous TRIM24 was immunoprecipitated as described above (TRIM24 - Proteintech #14208-1-AP). Samples were analyzed by immunoblotting.

### Luciferase reporter assays

Transient luciferase expression assays in HEK293T cells were performed as previously described (Dahabieh *et al*., 2011). Briefly, transfections were performed in 96 well plates seeded with 2×10^4^ HEK293T cells per well 24 hours prior to transfection. 10 ng of pGL3 reporter plasmid along with 10 ng of either pcDNA3.1+ (Invitrogen) or pcDNA-Tat and 100 ng expression vector was co-transfected. Luciferase activity was measured 24 hours post transfection. For Jurkat luciferase reporter assays, 1×10^5^ luciferase expressing Jurkat cells were plated with 100 µL media in 96-well plates. Luciferase activity was measured after the indicated time of treatment. Measurements were performed using Superlight™ luciferase reporter Gene Assay Kit (BioAssay Systems) as per the manufacturer’s instructions; 96 well plates were read in a VictorTM X3 Multilabel Plate Reader.

### Q-RT-PCR

RNA was extracted from Jurkat Tat KB001 cells following the indicated treatment using RNeasy Kit (Qiagen). RNA was analyzed using the Quant Studio 3 Real-Time PCR system (Applied Biosystems) using *Power* SYBR® Green RNA-to-CT™ 1-Step Kit (Thermo Fisher) as per the manufacturer’s instructions. Primers for analysis of HIV transcripts were as follows: Initiation, Fwd 5’ GTTAGACCAGATCTGAGCCT, Rev 5’ GTGGGTTCCCTAGTTAGCCA; Proximal, Fwd 5’ TGGGAGCTCTCTGGCTAACT, Rev 5’ TGCTAGAGATTTTCCACACTGA; Gag, Fwd 5’ CTAGAACGATTCGCAGTTAATCCT, Rev 5’ CTATCCTTTGATGCACACAATAGAG.

### Statistical analyses

Details of statistical analysis are indicated in Figure Legends. Mean is shown with standard deviations. Unpaired samples *t*-tests were performed with use of GraphPad Prism 9.0.0, and statistical significance is indicated at ^*^*P* < 0.05, ^**^*P* < 0.01, or ^***^*P* < 0.001.

## Acknowledgments

We thank Andy Johnson and Justin Wong of the UBC Flow Cytometry Facility for performing FACS analysis as well as for assistance with flow cytometry. This research was supported by program project grant F16-01210, from the Canadian Institutes of Health Research.

## Author Contributions

Malcolm T. performed the yeast-two hybrid screen that identified the TFII-I/ TRIM24 interaction. Dahabieh M. performed the experiments described in Figures 3 and EV4. Horvath R. M. performed all other experiments. Sadowski I. and Horvath R. M. wrote the paper.

## Legends to Expanded View Figures

**Figure EV1.**
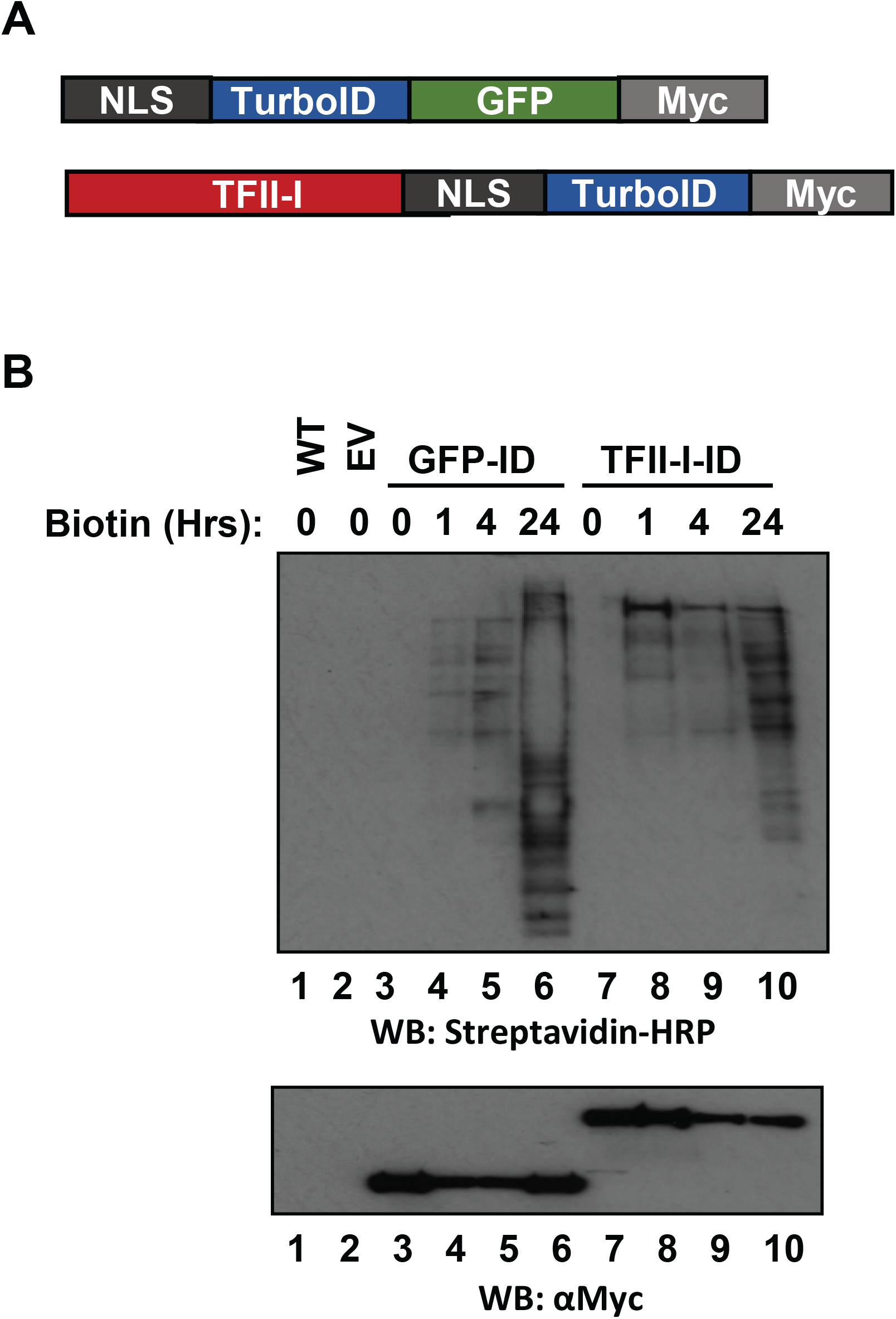
TurboID fusion proteins expressed in HEK293 cells. **Panel A:** Schematic representation of myc epitope tagged GFP (top) and TFII-I (bottom) TurboID fusions. **Panel B:** HEK293T cells (lane 1) were transfected with plasmids expressing GFP-TurboID (lanes 3-6), TFII-I-TurboID (lanes 7-10), or an empty vector control (EV, lane 2). Cells were incubated with 500 µM biotin for the indicated amount of time (lanes 3-10), when lysates were prepared and immunoblotted using Streptavidin-HRP (top) or anti-myc antibodies (bottom).

**Figure EV2.**
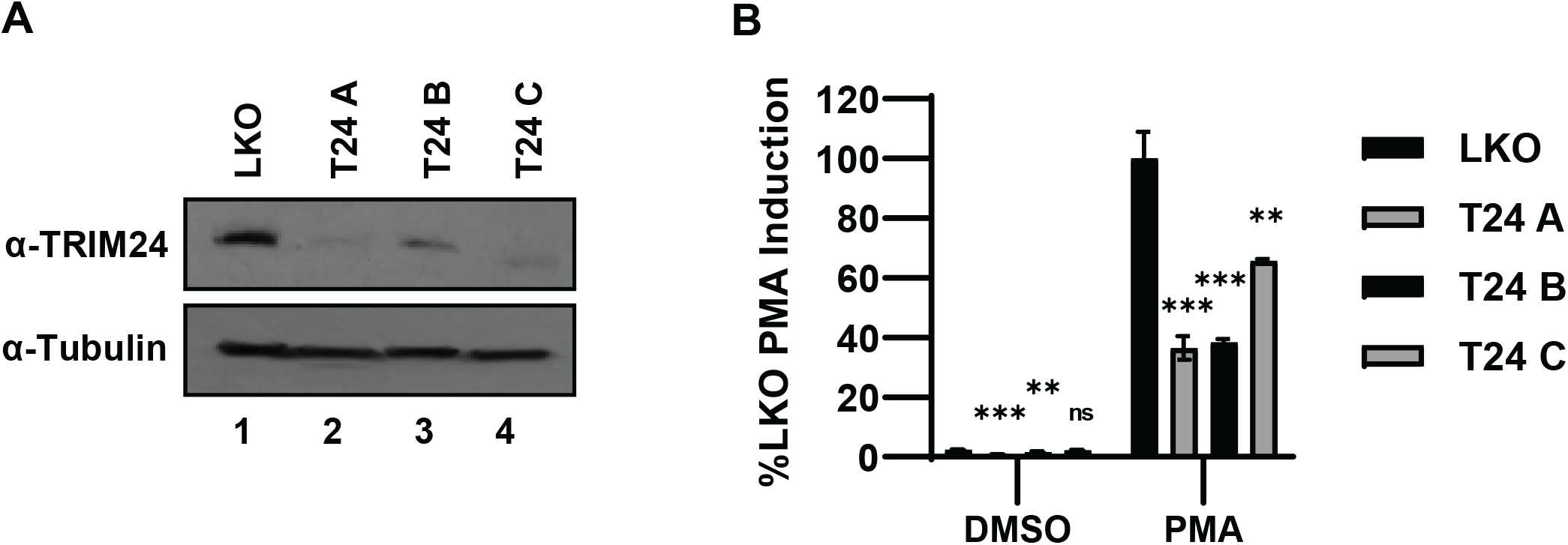
TRIM24 shRNA knockdown suppresses HIV-1 expression. **Panel A:** KB001 cells bearing an HIV-1-luciferase reporter provirus, were transduced with pLKO empty vector control (lane 1) or three shRNAs (A, B and C) targeting TRIM24 (lanes 2-4). Four days post infection and puromycin selection, lysates from transduced cells were analyzed by immunoblotting with antibodies against TRIM24 (top) or tubulin (bottom). **Panel B:** KB001 cells transduced with LKO vector control, or expressing TFII-I shRNAs were left untreated (DMSO) or stimulated with 20 nM PMA for 4 hours prior to measuring luciferase activity. Assays were performed in triplicate and error bars represent standard deviations.

**Figure EV3.**
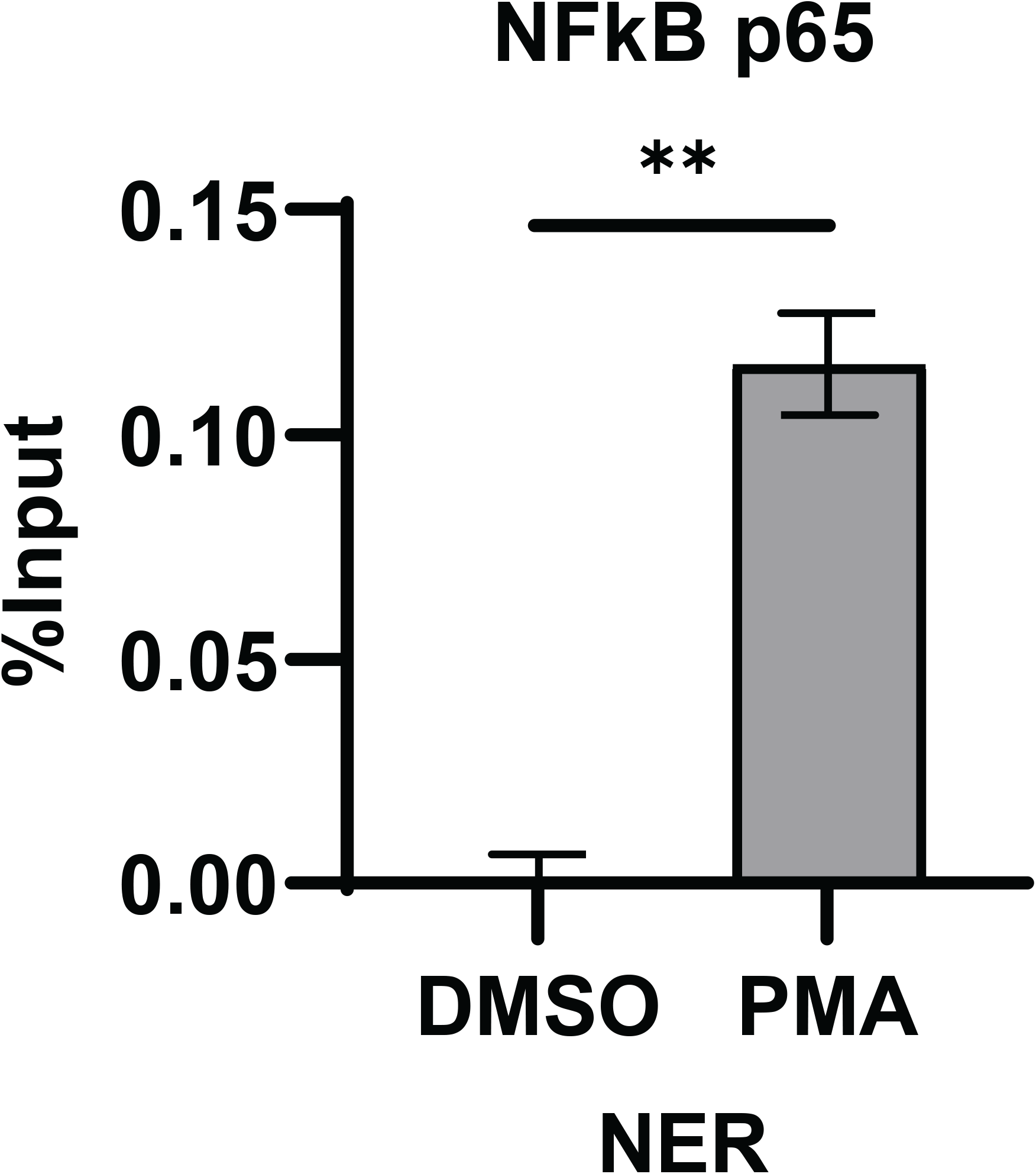
NFκB p65 is recruited to the LTR upon T cell activation. ChIP-Q PCR analysis with antibodies against p65 was performed with Jurkat Tat mdHIV clone 11 from untreated cells (DMSO) or stimulated with 50 nM PMA for 24 hrs. Results are an average of 2 independent measurements, and error bars represent standard deviations.

**Figure EV4.**
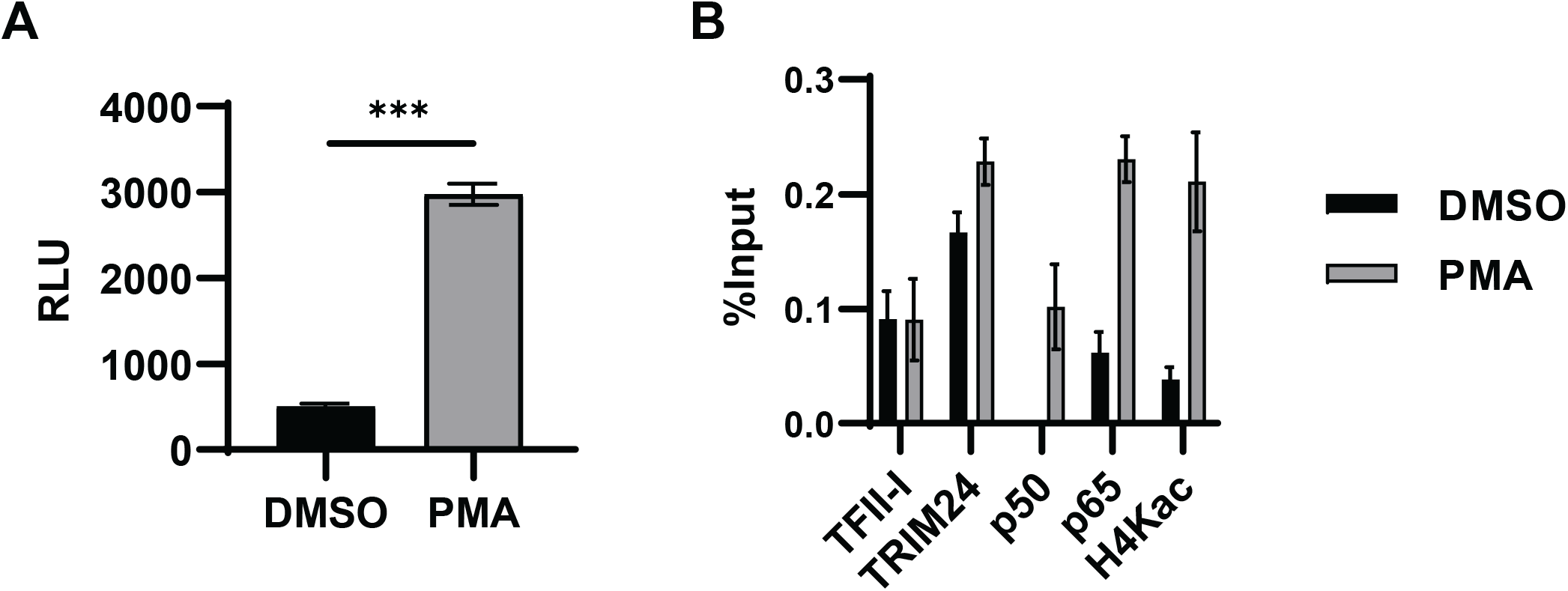
TRIM24 and TFII-I co-localize to the LTR. **Panel A:** TZM-bl cells were treated with DMSO or PMA (100 ng/mL) for 4 hrs prior to measurement of luciferase activity. Error bars represent standard deviations from triplicate experiments. **Panel B:** ChIP-qPCR with TZM-bl cells treated with DMSO or PMA was performed using the indicated antibodies. Results are an average of three measurements, and error bars represent standard deviations.

**Figure EV5.**
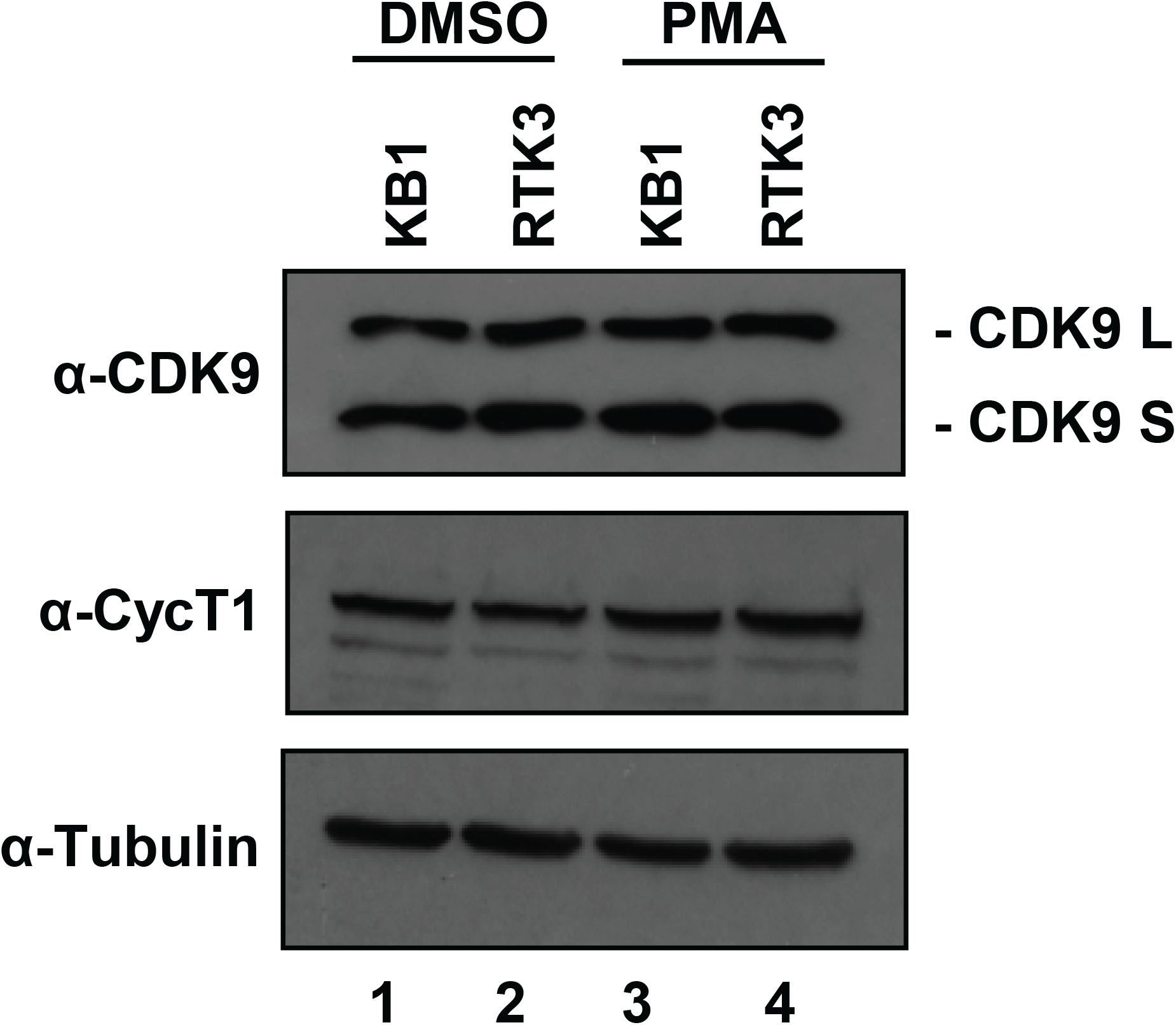
TRIM24 depletion does not alter CDK9 protein levels. KB001 or RTK3 *trim24* KO cells were left untreated (DMSO) (lanes 1-2) or treated with 20 nM PMA (lanes 3-4) for 4 hrs. Lysates were analyzed by immunoblotting with antibodies against CDK9, Cyclin T1, or tubulin as indicated.

**Figure EV6.**
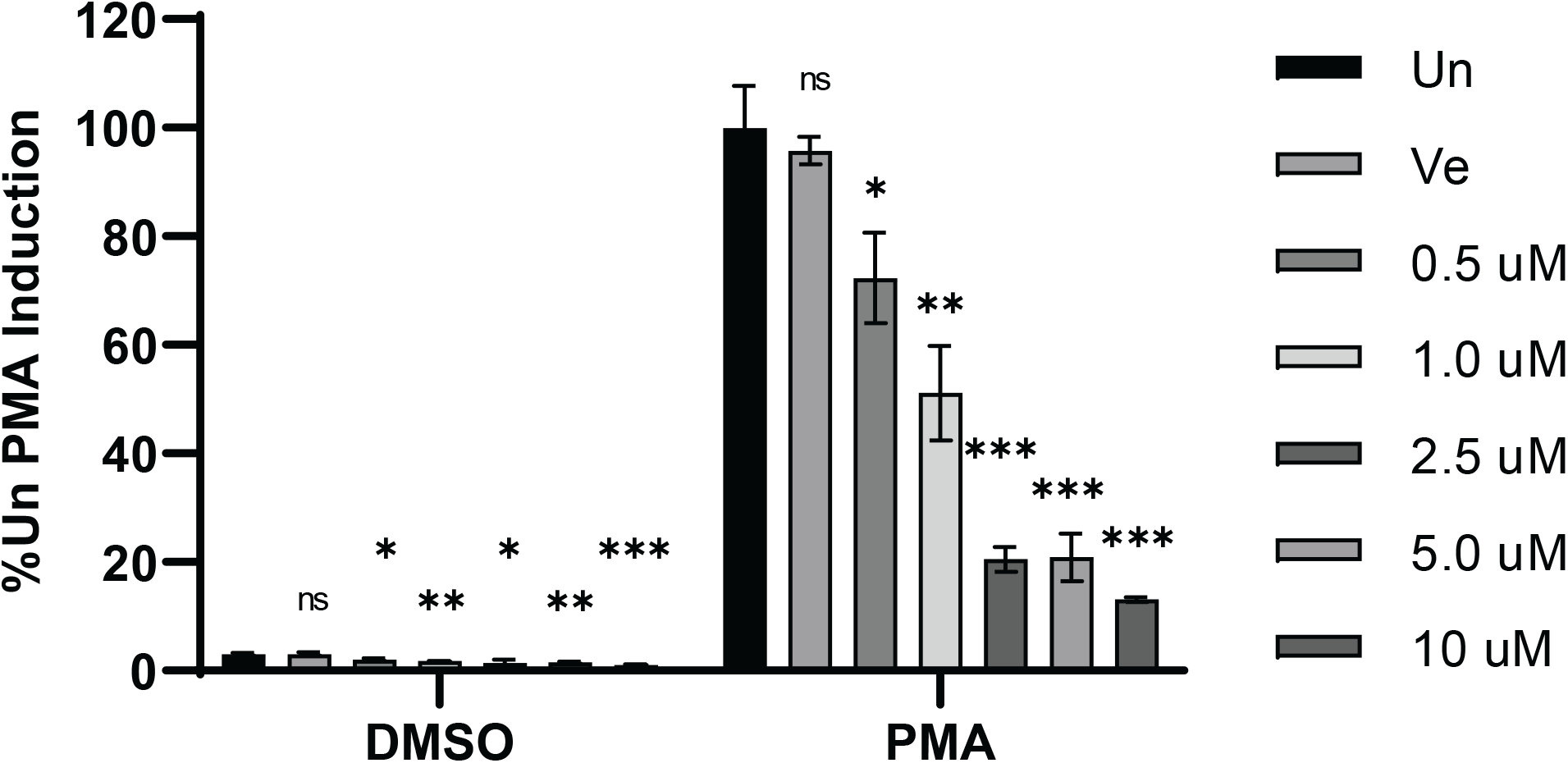
CDK9 kinase activity is necessary for induction of HIV-1 expression. Following 1 hr pre-treatment with the indicated concentration of CDK9 kinase inhibitor LDC67, KB001 cells were left untreated (DMSO) or incubated with 20 nM PMA for 4 hrs prior to luciferase measurement. Results are an average of three determinations, and error bars represent standard deviations.

## References

Allton K, Jain AK, Herz H-M, Tsai W-W, Jung SY, Qin J, Bergmann A, Johnson RL & Barton MC (2009) Trim24 targets endogenous p53 for degradation. Proc Natl Acad Sci U S A 106: 11612–11616

Appikonda S, Thakkar KN & Barton MC (2016) Regulation of gene expression in human cancers by TRIM24. Drug Discov Today Technol 19: 57–63

Appikonda S, Thakkar KN, Shah PK, Dent SYR, Andersen JN & Barton MC (2018) Cross-talk between chromatin acetylation and SUMOylation of tripartite motif-containing protein 24 (TRIM24) impacts cell adhesion. J Biol Chem 293: 7476–7485

Bell B & Sadowski I (1996) Ras-responsiveness of the HIV-1 LTR requires RBF-1 and RBF-2 binding sites. Oncogene 13: 2687–2697

Bernhard W, Barreto K, Raithatha S & Sadowski I (2013) An upstream YY1 binding site on the HIV-1 LTR contributes to latent infection. PLoS One 8: e77052

Bieniasz PD, Grdina TA, Bogerd HP & Cullen BR (1999) Recruitment of cyclin T1/P-TEFb to an HIV type 1 long terminal repeat promoter proximal RNA target is both necessary and sufficient for full activation of transcription. Proc Natl Acad Sci U S A 96: 7791–7796

Brooks DG, Arlen PA, Gao L, Kitchen CMR & Zack JA (2003) Identification of T cell-signaling pathways that stimulate latent HIV in primary cells. Proc Natl Acad Sci U S A 100: 12955–12960

Buratowski S (2009) Progression through the RNA polymerase II CTD cycle. Mol Cell 36: 541–546

Chen J, Malcolm T, Estable MC, Roeder RG & Sadowski I (2005) TFII-I regulates induction of chromosomally integrated human immunodeficiency virus type 1 long terminal repeat in cooperation with USF. J Virol 79: 4396–4406

Chun TW, Stuyver L, Mizell SB, Ehler LA, Mican JA, Baseler M, Lloyd AL, Nowak MA & Fauci AS (1997) Presence of an inducible HIV-1 latent reservoir during highly active antiretroviral therapy. Proc Natl Acad Sci U S A 94: 13193–13197

Corre S & Galibert M-D (2005) Upstream stimulating factors: highly versatile stress-responsive transcription factors. Pigment Cell Res 18: 337–348

Dahabieh MS, Ooms M, Malcolm T, Simon V & Sadowski I (2011) Identification and functional analysis of a second RBF-2 binding site within the HIV-1 promoter. Virology 418: 57–66

Durocher Y, Perret S & Kamen A (2002) High-level and high-throughput recombinant protein production by transient transfection of suspension-growing human 293-EBNA1 cells. Nucleic Acids Res 30: E9

Estable MC, Bell B, Merzouki A, Montaner JS, O’Shaughnessy MV & Sadowski IJ (1996) Human immunodeficiency virus type 1 long terminal repeat variants from 42 patients representing all stages of infection display a wide range of sequence polymorphism and transcription activity. J Virol 70: 4053–4062

Estable MC, Hirst M, Bell B, O’Shaughnessy MV & Sadowski I (1999) Purification of RBF-2, a transcription factor with specificity for the most conserved cis-element of naturally occurring HIV-1 LTRs. J Biomed Sci 6: 320–332

Finzi D, Hermankova M, Pierson T, Carruth LM, Buck C, Chaisson RE, Quinn TC, Chadwick K, Margolick J, Brookmeyer R, et al (1997) Identification of a reservoir for HIV-1 in patients on highly active antiretroviral therapy. Science 278: 1295–1300

Groner AC, Cato L, de Tribolet-Hardy J, Bernasocchi T, Janouskova H, Melchers D, Houtman R, Cato ACB, Tschopp P, Gu L, et al (2016) TRIM24 Is an Oncogenic Transcriptional Activator in Prostate Cancer. Cancer Cell 29: 846–858

Hashemi P, Barreto K, Bernhard W, Lomness A, Honson N, Pfeifer TA, Harrigan PR & Sadowski I (2018) Compounds producing an effective combinatorial regimen for disruption of HIV-1 latency. EMBO Mol Med 10: 160–174

Hatakeyama S (2017) TRIM Family Proteins: Roles in Autophagy, Immunity, and Carcinogenesis. Trends Biochem Sci 42: 297–311

Herquel B, Ouararhni K, Khetchoumian K, Ignat M, Teletin M, Mark M, Béchade G, Van Dorsselaer A, Sanglier-Cianférani S, Hamiche A, et al (2011) Transcription cofactors TRIM24, TRIM28, and TRIM33 associate to form regulatory complexes that suppress murine hepatocellular carcinoma. Proc Natl Acad Sci U S A 108: 8212–8217

Hirst M, Ho C, Sabourin L, Rudnicki M, Penn L & Sadowski I (2001) A two-hybrid system for transactivator bait proteins. Proc Natl Acad Sci U S A 98: 8726–8731

Jain AK, Allton K, Duncan AD & Barton MC (2014) TRIM24 is a p53-induced E3-ubiquitin ligase that undergoes ATM-mediated phosphorylation and autodegradation during DNA damage. Mol Cell Biol 34: 2695–2709

Joos B, Fischer M, Kuster H, Pillai SK, Wong JK, Böni J, Hirschel B, Weber R, Trkola A, Günthard HF, et al (2008) HIV rebounds from latently infected cells, rather than from continuing low-level replication. Proc Natl Acad Sci U S A 105: 16725–16730

Kanno Y, Kure Y, Kobayashi S, Mizuno M, Tsuchiya Y, Yamashita N, Nemoto K & Inouye Y (2018) Tripartite Motif Containing 24 Acts as a Novel Coactivator of the Constitutive Active/Androstane Receptor. Drug Metab Dispos 46: 46–52

Karn J (2011) The molecular biology of HIV latency: breaking and restoring the Tat-dependent transcriptional circuit. Curr Opin HIV AIDS 6: 4–11

Kikuchi M, Okumura F, Tsukiyama T, Watanabe M, Miyajima N, Tanaka J, Imamura M & Hatakeyama S (2009) TRIM24 mediates ligand-dependent activation of androgen receptor and is repressed by a bromodomain-containing protein, BRD7, in prostate cancer cells. Biochim Biophys Acta 1793: 1828–1836

Kim DW & Cochran BH (2000) Extracellular signal-regulated kinase binds to TFII-I and regulates its activation of the c-fos promoter. Mol Cell Biol 20: 1140–1148

Laskey SB & Siliciano RF (2014) A mechanistic theory to explain the efficacy of antiretroviral therapy. Nat Rev Microbiol 12: 772–780

Lv D, Li Y, Zhang W, Alvarez AA, Song L, Tang J, Gao W-Q, Hu B, Cheng S-Y & Feng H (2017) TRIM24 is an oncogenic transcriptional co-activator of STAT3 in glioblastoma. Nat Commun 8: 1454

Malcolm T, Chen J, Chang C & Sadowski I (2007) Induction of chromosomally integrated HIV-1 LTR requires RBF-2 (USF/TFII-I) and Ras/MAPK signaling. Virus Genes 35: 215–223

Malcolm T, Kam J, Pour PS & Sadowski I (2008) Specific interaction of TFII-I with an upstream element on the HIV-1 LTR regulates induction of latent provirus. FEBS Lett 582: 3903–3908

Mbonye U & Karn J (2014) Transcriptional control of HIV latency: cellular signaling pathways, epigenetics, happenstance and the hope for a cure. Virology 454–455: 328–339

Mbonye UR, Gokulrangan G, Datt M, Dobrowolski C, Cooper M, Chance MR & Karn J (2013) Phosphorylation of CDK9 at Ser175 enhances HIV transcription and is a marker of activated P-TEFb in CD4(+) T lymphocytes. PLoS Pathog 9: e1003338

Mbonye U, Wang B, Gokulrangan G, Shi W, Yang S & Karn J (2018) Cyclin-dependent kinase 7 (CDK7)-mediated phosphorylation of the CDK9 activation loop promotes P-TEFb assembly with Tat and proviral HIV reactivation. J Biol Chem 293: 10009–10025

McAvera RM & Crawford LJ (2020) TIF1 Proteins in Genome Stability and Cancer. Cancers (Basel) 12: E2094

Pathiraja TN, Thakkar KN, Jiang S, Stratton S, Liu Z, Gagea M, Shi X, Shah PK, Phan L, Lee M-H, et al (2015) TRIM24 links glucose metabolism with transformation of human mammary epithelial cells. Oncogene 34: 2836–2845

Pereira LA, Bentley K, Peeters A, Churchill MJ & Deacon NJ (2000) A compilation of cellular transcription factor interactions with the HIV-1 LTR promoter. Nucleic Acids Res 28: 663–668

Roy AL (2001) Biochemistry and biology of the inducible multifunctional transcription factor TFII-I. Gene 274: 1–13

Roy AL, Meisterernst M, Pognonec P & Roeder RG (1991) Cooperative interaction of an initiator-binding transcription initiation factor and the helix-loop-helix activator USF. Nature 354: 245–248

Sadowski I & Hashemi FB (2019) Strategies to eradicate HIV from infected patients: elimination of latent provirus reservoirs. Cell Mol Life Sci 76: 3583–3600

Sadowski I, Lourenco P & Malcolm T (2008) Factors controlling chromatin organization and nucleosome positioning for establishment and maintenance of HIV latency. Curr HIV Res 6: 286–295

Thénot S, Henriquet C, Rochefort H & Cavaillès V (1997) Differential interaction of nuclear receptors with the putative human transcriptional coactivator hTIF1. J Biol Chem 272: 12062–12068

Tisserand J, Khetchoumian K, Thibault C, Dembélé D, Chambon P & Losson R (2011) Tripartite motif 24 (Trim24/Tif1α) tumor suppressor protein is a novel negative regulator of interferon (IFN)/signal transducers and activators of transcription (STAT) signaling pathway acting through retinoic acid receptor α (Rarα) inhibition. J Biol Chem 286: 33369–33379

Tsai W-W, Wang Z, Yiu TT, Akdemir KC, Xia W, Winter S, Tsai C-Y, Shi X, Schwarzer D, Plunkett W, et al (2010) TRIM24 links a non-canonical histone signature to breast cancer. Nature 468: 927–932

Verdin E, Paras P & Van Lint C (1993) Chromatin disruption in the promoter of human immunodeficiency virus type 1 during transcriptional activation. EMBO J 12: 3249–3259

Vullhorst D & Buonanno A (2005) Multiple GTF2I-like repeats of general transcription factor 3 exhibit DNA binding properties. Evidence for a common origin as a sequence-specific DNA interaction module. J Biol Chem 280: 31722–31731

Wen Y-D, Cress WD, Roy AL & Seto E (2003) Histone deacetylase 3 binds to and regulates the multifunctional transcription factor TFII-I. J Biol Chem 278: 1841–1847

Wong JK, Hezareh M, Günthard HF, Havlir DV, Ignacio CC, Spina CA & Richman DD (1997) Recovery of replication-competent HIV despite prolonged suppression of plasma viremia. Science 278: 1291–1295

Yu T, Gan S, Zhu Q, Dai D, Li N, Wang H, Chen X, Hou D, Wang Y, Pan Q, et al (2019) Modulation of M2 macrophage polarization by the crosstalk between Stat6 and Trim24. Nat Commun 10: 4353

Yukl SA, Kaiser P, Kim P, Telwatte S, Joshi SK, Vu M, Lampiris H & Wong JK (2018) HIV latency in isolated patient CD4+ T cells may be due to blocks in HIV transcriptional elongation, completion, and splicing. Sci Transl Med 10: eaap9927

Zhu Q, Yu T, Gan S, Wang Y, Pei Y, Zhao Q, Pei S, Hao S, Yuan J, Xu J, et al (2020) TRIM24 facilitates antiviral immunity through mediating K63-linked TRAF3 ubiquitination. J Exp Med 217: e20192083

